# Deep Semantic Protein Representation for Annotation, Discovery, and Engineering

**DOI:** 10.1101/365965

**Authors:** Ariel S Schwartz, Gregory J Hannum, Zach R Dwiel, Michael E Smoot, Ana R Grant, Jason M Knight, Scott A Becker, Jonathan R Eads, Matthew C LaFave, Harini Eavani, Yinyin Liu, Arjun K Bansal, Toby H Richardson

**Affiliations:** Synthetic Genomics Inc., La Jolla, California, USA; Intel AI Lab, San Diego, California, USA

## Abstract

Computational assignment of function to proteins with no known homologs is still an unsolved problem. We have created a novel, function-based approach to protein annotation and discovery called D-SPACE (Deep Semantic Protein Annotation Classification and Exploration), comprised of a multi-task, multi-label deep neural network trained on over 70 million proteins. Distinct from homology and motif-based methods, D-SPACE encodes proteins in high-dimensional representations (embeddings), allowing the accurate assignment of over 180,000 labels for 13 distinct tasks. The embedding representation enables fast searches for functionally related proteins, including homologs undetectable by traditional approaches. D-SPACE annotates all 109 million proteins in UniProt in under 35 hours on a single computer and searches the entirety of these in seconds. D-SPACE further quantifies the relative functional effect of mutations, facilitating rapid in silico mutagenesis for protein engineering applications. D-SPACE incorporates protein annotation, search, and other exploratory efforts into a single cohesive model.

---

We are witnessing an explosion in biological sequence information^1^ stored in databases such as GenBank^2^ and UniProt^3^, largely driven by massive improvements in sequencing technology and resulting in the availability of tens of thousands of genomes and metagenomes. The utility of each genome depends on high quality annotation and the tools available to analyze it, shifting the bottleneck for genomics-based scientific discovery from sequencing capacity to analysis. While genome annotation has evolved from a laborious manual process to a largely automated endeavor with a proliferation of sophisticated tools and pipelines, many proteins, including some essential for life^4^, still have no known function. To date, the ability to reliably predict protein functions directly from amino-acid sequences alone remains a major biological challenge^5^.

Since the function of a protein is a direct consequence of its amino acid sequence, similarity between primary sequence has been used to systematically infer function for more than 25 years^6^. While generally useful, a simple similarity measure is often insufficient to confidently assign protein function. Highly divergent natural sequences sometimes have similar functions, and even single amino acid changes can completely eliminate the function of a protein. In addition, many proteins have no known homologs. More complex statistical models such as profile hidden Markov models HMMs (pHMMs) have been developed to address these challenges^7–12^, and while highly beneficial, such approaches often lack generalizability. Each pHMM model is typically trained on hand-curated aligned sequences of a given protein family without leveraging information from other families or annotation types, which results in very high specificity for the trained family without the ability to detect functionally-related but sequence-divergent proteins. These models are also computationally intensive to run at scale, which can be challenging for annotating large protein databases or metagenomes. More recently, databases such as InterPro^13^ and PROSITE^14^ have aimed at integrating genomic features such as domain and functional site information from diverse databases into a single searchable user interface. To our knowledge there is not one consistent model that is able to learn from all of this information to predict all features.

Making the full connection between sequence and function requires a representation of a protein that is closely associated with broad range of functional properties, which is now possible due to advances in high-level representations from the field of deep-learning. Several groups have deployed machine learning methods to identify specific functions of proteins^15–19^, however there have been only limited attempts to build a single classifier that assigns multiple features to a given peptide sequence^20–23^.

Our work builds on these ideas with knowledge integration at massive scale, both in the number of proteins trained on and in the annotations used. By building the deep and comprehensive multi-task model D-SPACE, we created a high-dimensional protein representation (‘embedding’) which can be used to improve many sequence-based informatics tasks. The D-SPACE model can annotate proteins extremely quickly and combines the most relevant features of more than 180,000 smaller models.

The protein embedding space provides a content-rich representation which enables the determination of many aspects of a protein structure and function from a simple Euclidean coordinate. This property allows for ultra-fast search for functionally similar proteins, even for those with highly divergent sequence. We show evidence that this representation generalizes to proteins on which the model was not trained. This approach represents a foundation for integrating all existing and future knowledge of protein structure and function into a cohesive and generalizable model. A demo of the D-SPACE annotation and search features is available at https://dspace.bio.

## RESULTS

### D-SPACE model training on 70 million unique proteins

We trained our model using protein sequences from UniProtKB (Swiss-Prot and TrEMBL), filtered to exclude any duplicate proteins that are not the representative member of a UniRef100 cluster^3, 24^. We also included or constructed annotations from 13 sources (**Table 1**, **Methods**). In total we assembled more than 90 million records of which 80% were used as the training set, 10% were used for validating the model training, and another 10% were held out as the test set for final evaluation of the model’s performance. Our model is based on a deep convolutional neural network architecture with more than 100 million trainable parameters (**Fig. 1a**, **Supplementary Fig. 1**). Part of this model is a convergent affine ‘embedding’ layer consisting of 256 floating-point values, from which all classification outputs are derived (**Fig. 1b**). As an additional output, the model includes an autoencoder to compress the 256-dimension protein embedding to a non-linear three-dimensional representation (**Fig. 1c, 1d**).

**Table 1:**
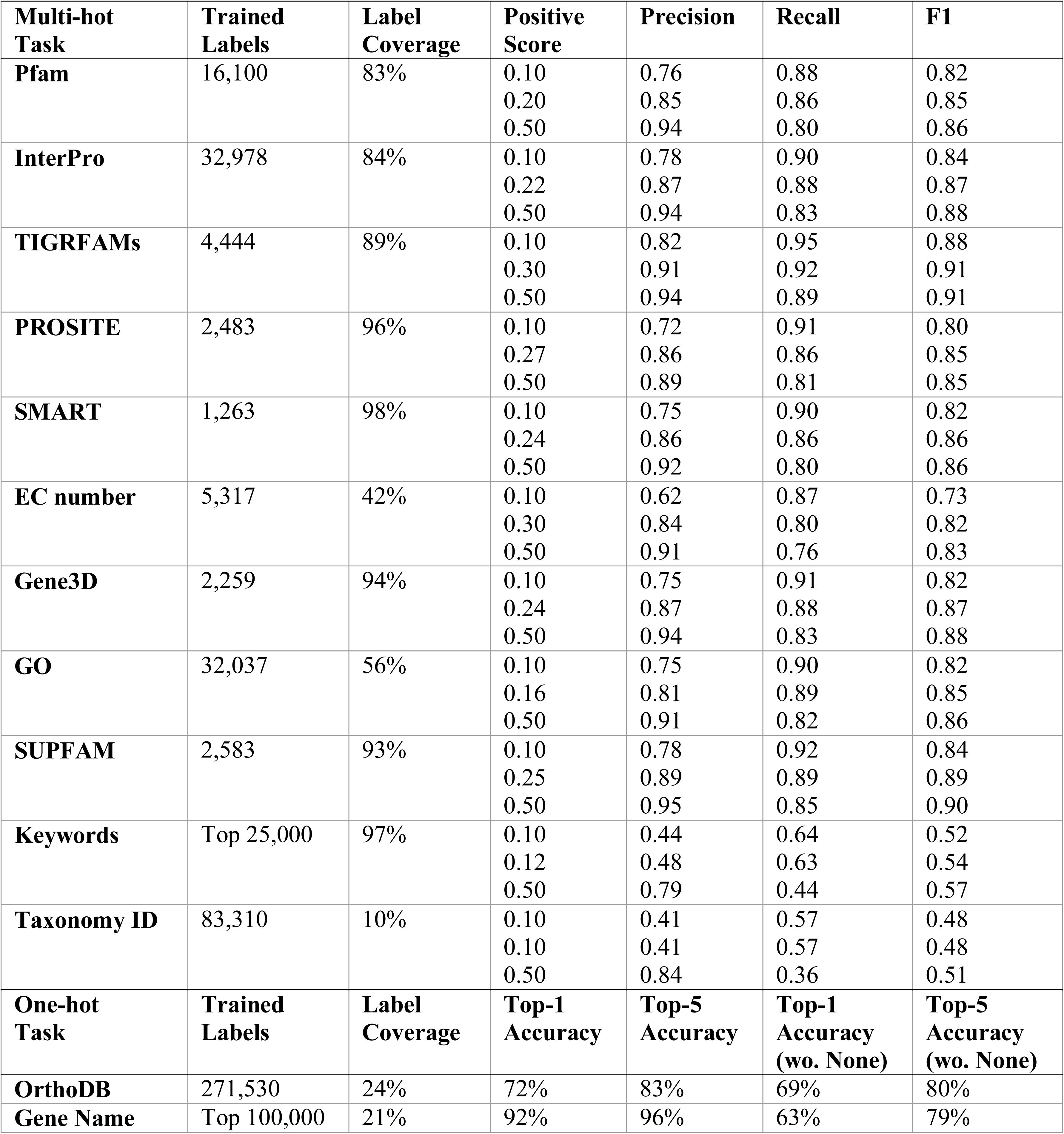
List and performance of D-SPACE annotation tasks. Label Coverage is a measure of the representation of positive calls across labels in the test set. It is calculated as the number of unique labels with a positive call (Top-5 for one-hot, Score > 0.1 for multi-hot) divided by the number of unique true labels in the test set. Accuracy measures for one-hot labels are taken as either Top-1 or Top-5, both including and not including the default label ‘None’. Performance measures are reported for positive scores at the permissive 0.1 value, the ‘optimal’ value (**Methods**), and the default 0.5 value.

**Figure 1.**
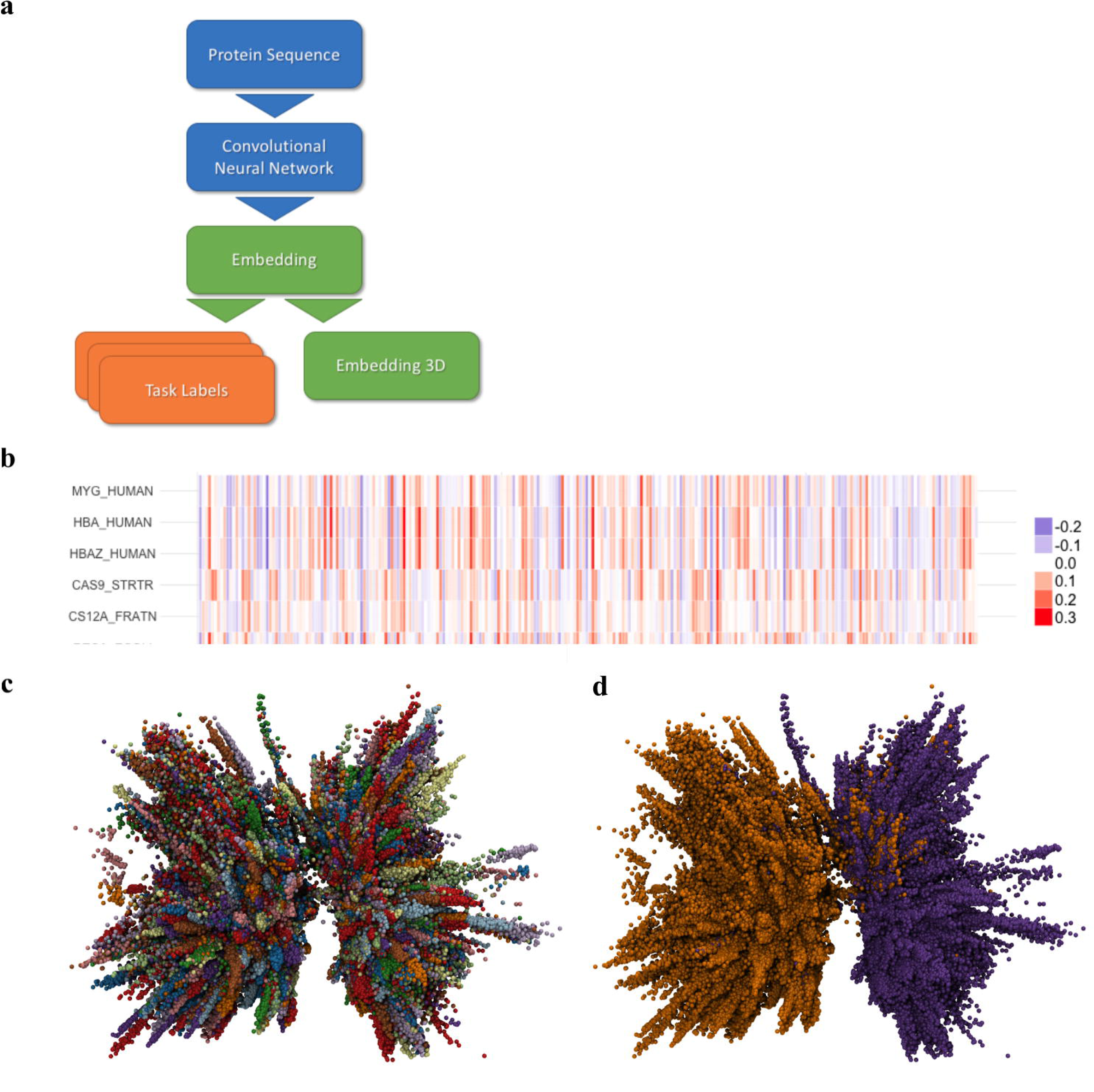
D-SPACE encodes a high-dimensional representation of a protein. (**a**) A broad schematic of the D-SPACE model. Protein sequences are encoded with a convolutional neural network to produce a 256-dimenstional ‘embedding’ representation. This can be used to infer labels for multiple tasks or reduced to three dimensions for visualization. (**b**) Example embedding vectors for: homologous proteins with different degrees of sequence divergence (human myoglobin, hemoglobin and fetal hemoglobin); functionally related but sequence divergent proteins (Cas9 and Cas12a); and an unrelated protein (bacterial RecA). The colors represent the numeric values of the embedding. (**c**) The three-dimensional representation of more than 460,000 proteins from the Swiss-Prot database, colored by the true OrthoDB cluster label. Proteins in the same sequence cluster were grouped together. (**d**) The same three-dimensional representation, colored by sequence length. The model finds clear functional separation between small (<300 aa, purple) and large (>=300 aa, orange) proteins.

The model was trained for nine days and approximately four full passes over the training set until the validation loss was relatively stable (**Methods**). The training step with lowest validation loss was saved and used for inference and further analysis. We then annotated the entire dataset, including the training, validation, and test sets in less than 35 hours using a single computer. There was little evidence of overfitting (training / validation loss = 0.9). We also found model performance to be insensitive to most hyperparameter changes and that many similar network architectures could produce comparable results.

### D-SPACE accurately annotates protein sequences

The D-SPACE model performed well on all 13 of the classification tasks in the previously unseen test set (**Table 1, Methods**). Notably, the D-SPACE model tended to predict fewer class labels than might be expected (‘Coverage’ column in **Table 1**). This was largely due to the model’s ability to recognize which labels it could confidently call and which were best omitted. The highest performing class labels tended to be the most common, but this was not always the case (**Supplementary Fig. 2**). While further development could improve performance, perfect recapitulation of existing tools was not our goal. We found many examples where ‘incorrect’ assignments were appropriate when investigated manually. One group of these examples were proteins with UniProt records which had not yet been fully annotated in the training or test sets at the time of data download but were subsequently updated in later versions of UniProt. The fully updated annotations generally matched D-SPACE’s high confidence predictions. One example is UniProt A0A2I1C1K8, which was unannotated in our data set, but D-SPACE assigned it to the OrthoDB cluster ‘Cytochrome P450’ (score = 0.99) and suggested the keywords ‘4-monooxygenase’ (score = 0.18) and ‘pisatin’ (score = 0.14). UniProt now annotates this protein as ‘Putative cytochrome P450 pisatin demethylase’. That D-SPACE learned to give such cases appropriate annotations rather than predict their absence is a sign of useful generalization.

**Figure 2.**
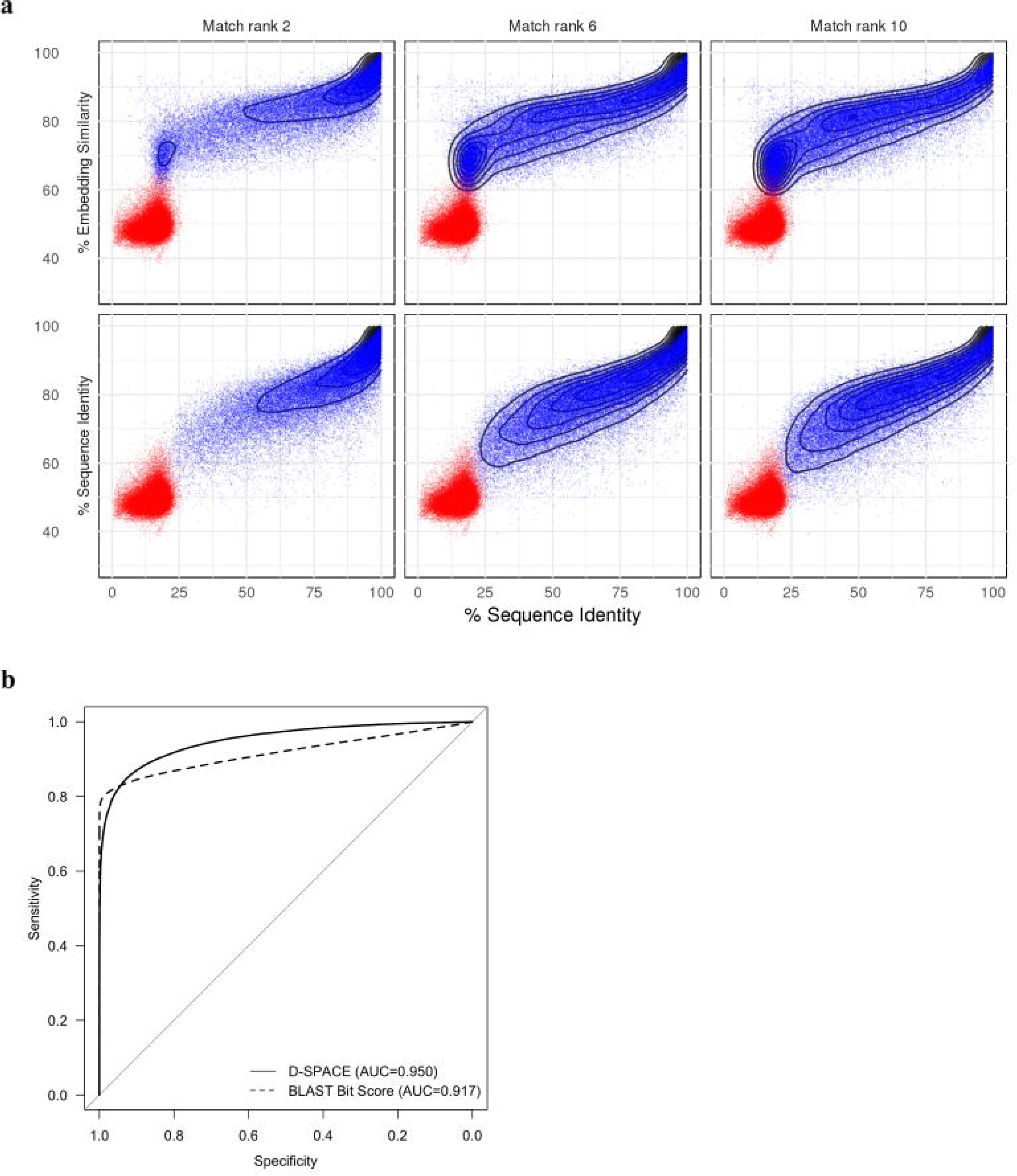
Comparison of sequence and embedding similarity for protein search. **(a**) Distribution of sequence and embedding similarity measures for related protein pairs (blue) and random pairs (orange). Related proteins are based on the 2^nd^, 6^th^, and 10^th^, best match using embedding similarity (top row) or sequence identity (bottom row). Proteins with high sequence identity (x-axis) were scored high with both approaches, but only embedding similarity tended to find matches below 25% sequence identity. **(b**) Comparison of sensitivity and specificity tradeoffs between sequence and embedding similarity measures for the Saripella *et al*. ortholog benchmark. Sequence similarity was slightly more sensitive at a high specificity threshold, but D-SPACE outperformed overall.

To assist in visualization of D-SPACE’s organization of the functional protein space, we included a three-dimensional (3D) embedding autoencoder in the model. The autoencoder was successful in reducing the dimensionality of the functional embedding space from 256 to 3 dimensions with only modest loss of information (mean-squared error = 0.001). The autoencoder also had a strong regularization effect, reducing the mean variance of the embedding layer by more than 75-fold as compared to a model trained without it. Visualizing the 3D space gives a partial view of the richness encoded by the model (**Fig. 1c, 1d**). As expected, similar sequences were found to be grouped closely in this representation.

### Protein functional embedding space enables fast and sensitive search and discovery

The key to the D-SPACE model’s success is its ability to map any protein sequence into a defined coordinate in a common 256-dimensional functional embedding space. In this embedding space, the distance between two proteins is determined by functional similarity rather than sequence similarity alone. Previous work has shown that remote homology searches can be accelerated when operating in a compressed embedding space learned from known sequence similarities^25–28^. D-SPACE generalizes this concept by learning a functional-based embedding space through multi-task^29^, multi-label training which also enables discovery of novel protein relationships beyond sequence homology.

An additional advantage of searching over the embedding space rather than sequence space is the ability to utilize advances in general case k-nearest neighbors (KNN) search tools and algorithms^30, 31^. In our implementation, a query against the full UniProtKB database (over 109 million proteins) returns the 100 nearest neighbors in less than 5 seconds, compared to several minutes with a standard BLAST installation. To evaluate the sensitivity of the embedding search results, we queried every protein in Swiss-Prot against the other Swiss-Prot proteins (552,000) using a sequence similarity-based approach and D-SPACE’s embedding search. Random protein pairs, which are not likely to be functionally related, tend to have sequence identity lower than 25% and embedding similarity lower than 0.55. There was a significant correlation between sequence identity and embedding similarity (R = 0.84, p-value < 2.2e-16) and both methods ranked highly similar proteins to the query sequence as their top results (**Fig. 2a**). However, while sequence-based methods could only detect hits with sequence identity >25%, embedding search was capable of identifying many functionally related hits below this level, in the ‘twilight-zone’ of sequence identity^32^. On the other hand, there were a few protein pairs identifiable by sequence identity-based methods with embedding similarity below the significant level of 0.55. In the cases we inspected, these potentially missed pairs were due to having only a partial local similarity between two proteins that shared domain similarity but had an overall different structure and function.

To verify that the additional D-SPACE matches are functionally relevant, we compared D-SPACE embedding similarity to sequence similarity using an established benchmark for detecting orthologous protein pairs developed by Saripella *et al*.^6, 33^. While sequence similarity had a slightly higher sensitivity for top hits (0.8 vs. 0.7 at 99% specificity), embedding similarity was more sensitive for other matches (0.92 vs 0.87 at 80% specificity) and had higher overall performance (AUC 0.95 vs. 0.92) (**Fig. 2b**).

### Functional embedding search enables novel protein discovery

As an example of the ability of the functional embedding search to extend into and beyond sequence similarity alone and into the twilight-zone of sequence homology, we searched UniProtKB using the rare Class 2 Type V CRISPR effector protein Cas12b (T0D7A2) with both BLAST and D-SPACE. The BLAST search ran for 3 minutes (using UniProt’s web interface) returning 21 significant hits (E-value < 0.1; excluding the self-hit to the query), of which only 7 aligned globally to the query sequence. All 21 hits are currently annotated as ‘Uncharacterized protein’ in UniProt, so they provided little immediate value in assigning function to the query sequence beyond the confirmation that they might belong to a related protein family. We also ran PSI-Blast and JackHMMER, which produced similar results^34, 35^. Using the embedding similarity search, D-SPACE was able to query Swiss-Prot and TrEMBL (returning the top 50 hits from each database) in less than 5 seconds, with all 100 hits having >0.59 embedding similarity to the query. In contrast to the BLAST results, D-SPACE’s results were enriched with currently annotated CRISPR-associated endonuclease and endoribonuclease proteins, including nine hits to Cas9 proteins, three hits to Cas13a proteins, and two hits to Cas12a (Cpf1) proteins, all with sequence identity <22% to the query sequence. Performing the same search with Cas12a (A0Q7Q2) as the query yielded similar results. Inspecting the three-dimensional representation of the embedding space revealed that Cas9 and Cas12a proteins tended to be placed close to each other (**Fig. 3a**). Since many Cas9 and some Cas12a were included in the original training set, one might assume that the model learned to associate them based on shared feature labels, such as keywords and Gene Ontology (GO) terms. To test this hypothesis, we trained a second D-SPACE model excluding all annotated Cas12a proteins from the training set and inspected the resulting embedding space (**Fig. 3b**). The results show that even when not presented with examples of Cas12a sequences during training, D-SPACE’s model was able to associate Cas12a with other Class 2 CRISPR effector proteins during inference, demonstrating the generalizability of D-SPACE’s predictions and its ability to identify novel proteins without relying on sequence similarity.

**Figure 3.**
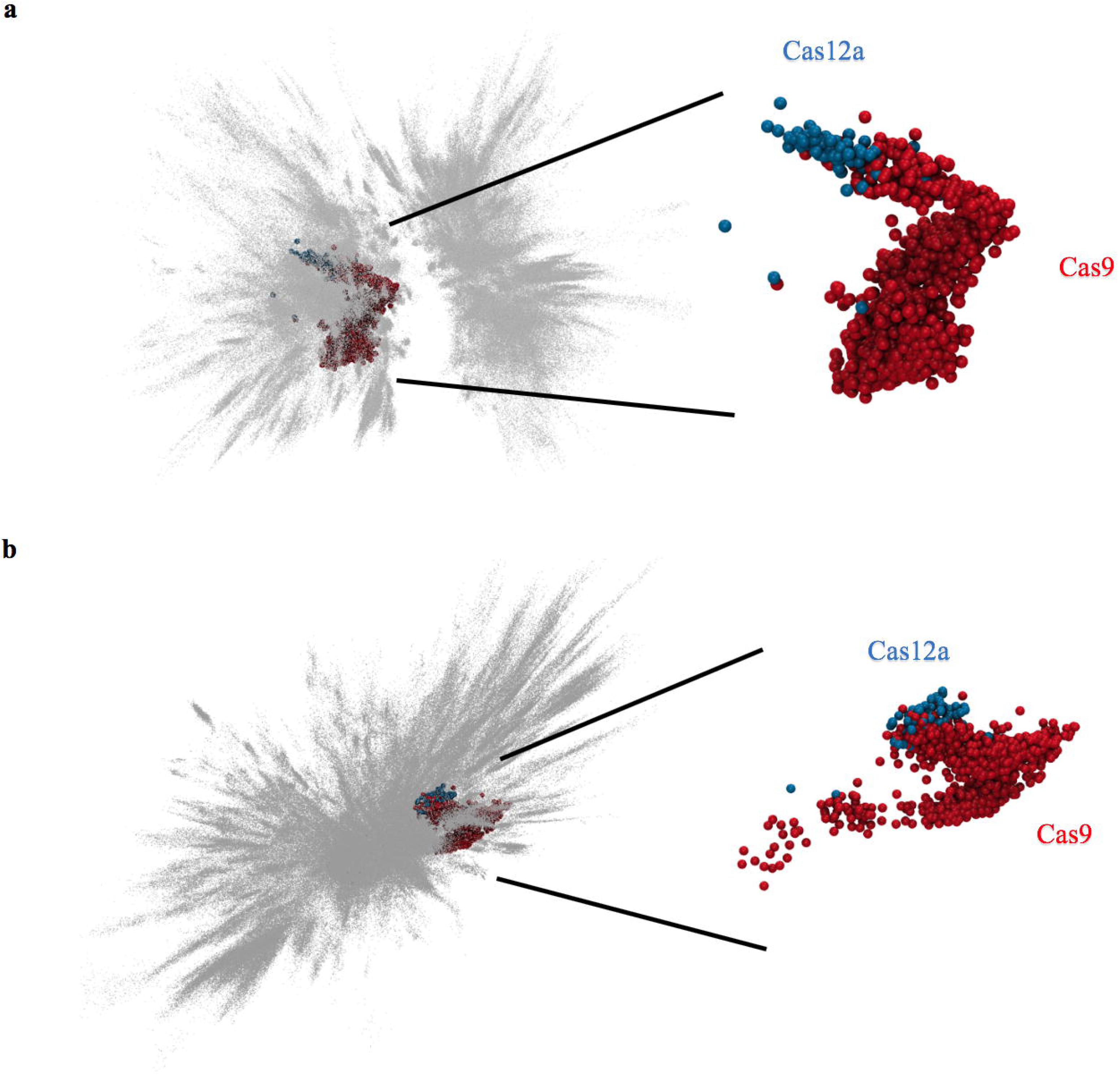
Cas9 and Cas12a comparison in D-SPACE. **(a**) Cas9 (red) and Cas12a (blue) proteins were mapped to the three-dimensional embedding space with Swiss-Prot proteins shown in gray. Despite their low sequence similarity, the proteins tended to cluster together. (**b**) The process was repeated for a D-SPACE model trained without any Cas12a examples. These proteins were still represented near each other, indicating functional similarity.

### Combining D-SPACE annotation and search for protein annotation

A recent study by Price *et al*. described new experimental approaches to identifying gene function for 33 bacterial organisms^36^. We used D-SPACE to annotate these same genomes computationally in less than an hour. The Price *et al.* work resulted in a non-trivial (e.g. not ‘hypothetical’) descriptions for 75% of the 156,184 proteins (**Methods**). D-SPACE produced a rich multi-label output for every protein. Of these, 84% were assigned an InterPro label (score > 0.22), 73% had a non-trivial top OrthoDB cluster (top result not ‘None’, score > 0.1), and the proteins overall averaged 18 GO terms each (score > 0.16). The search capabilities of D-SPACE were also used to annotate the proteins by finding functional homologs. To demonstrate this, we searched each of the bacterial proteins against Swiss-Prot. A total of 82% of proteins had a strong functional homolog with a non-trivial description (top result and score > 0.55). In all, 95% of proteins were assigned either an InterPro label, a non-trivial OrthoDB cluster, or a non-trivial functional homolog. This demonstrates a dramatic leap in our capability to assign putative functions to currently unknown proteins (75% to 95% of the dataset).

### Semantic embedding profiles for protein discovery

Functional embedding search requires only a coordinate in the high-dimensional embedding space as a query, and therefore does not rely on protein sequence information as its input. In addition to the enhanced performance of D-SPACE for the basic case of single protein query searches compared to sequence-based methods, advanced profile-based searches can be supported by D-SPACE by constructing novel semantic embedding profiles and using them as input queries. One such approach is to combine multiple embedding vectors using simple algebraic operations.

A common use case in traditional protein discovery workflows involves construction of a pHMM for a given protein family, which can then be used to search over protein sequence databases. While powerful, they can only reliably be constructed using proteins with similar domain structure and with a detectable level of sequence similarity. In contrast, functional embedding profiles can be constructed for any set of input embedding vectors, by calculating a profile embedding vector using simple aggregation functions (e.g. mean or median), resulting in a new coordinate in the embedding space that can be used as a query. In addition, embedding profiles can be combined to enable powerful semantic searches even when their underlying protein sequences have no detectable sequence similarity.

As an example of embedding profile search we calculated the mean embedding profile of all proteins in Swiss-Prot (‘p0’), as well as mean embedding profiles of two Type V DNA effector Cas12a proteins (‘p1’) and seven Type VI RNA effector Cas13a proteins (‘p2’). We then combined the Cas12a and Cas13a profiles into a joint profile ‘p12’ (p12 = p1 + p2 − p0). Searching with the joint profile against UniProtKB returned 59 hits (top 50 each from Swiss-Prot and TrEMBL filtered by similarity score > 0.55). Of these, 39 had a known function, including 25 Cas12a proteins, 8 Cas13a proteins, 3 Cas9 proteins, and 3 proteins with unrelated annotations. This demonstrates the ability to perform relevant searches using profiles from proteins with no sequence similarity.

### D-SPACE enables in silico protein mutagenesis analysis

We used the generalizability of the D-SPACE model to test in silico mutagenesis experiments, where each amino acid in a protein is individually replaced with all possible alternatives. Each mutated sequence was run through the model, and the resulting embedding vectors were compared to the original protein to reveal each mutation’s impact on overall protein function (**Fig. 4**). This processed identified signatures that correspond to key protein features. For one example, RecA, the signature coincided with the well-known NTP binding P-loop motif ^37^. In another example, Tpo1, the signature coincided with known transmembrane regions. These are exceptional findings given that no positional information was used in the model training process. D-SPACE was capable of identifying the relevant features on its own and can provide insight to specific modifications which are likely to impact function.

**Figure 4.**
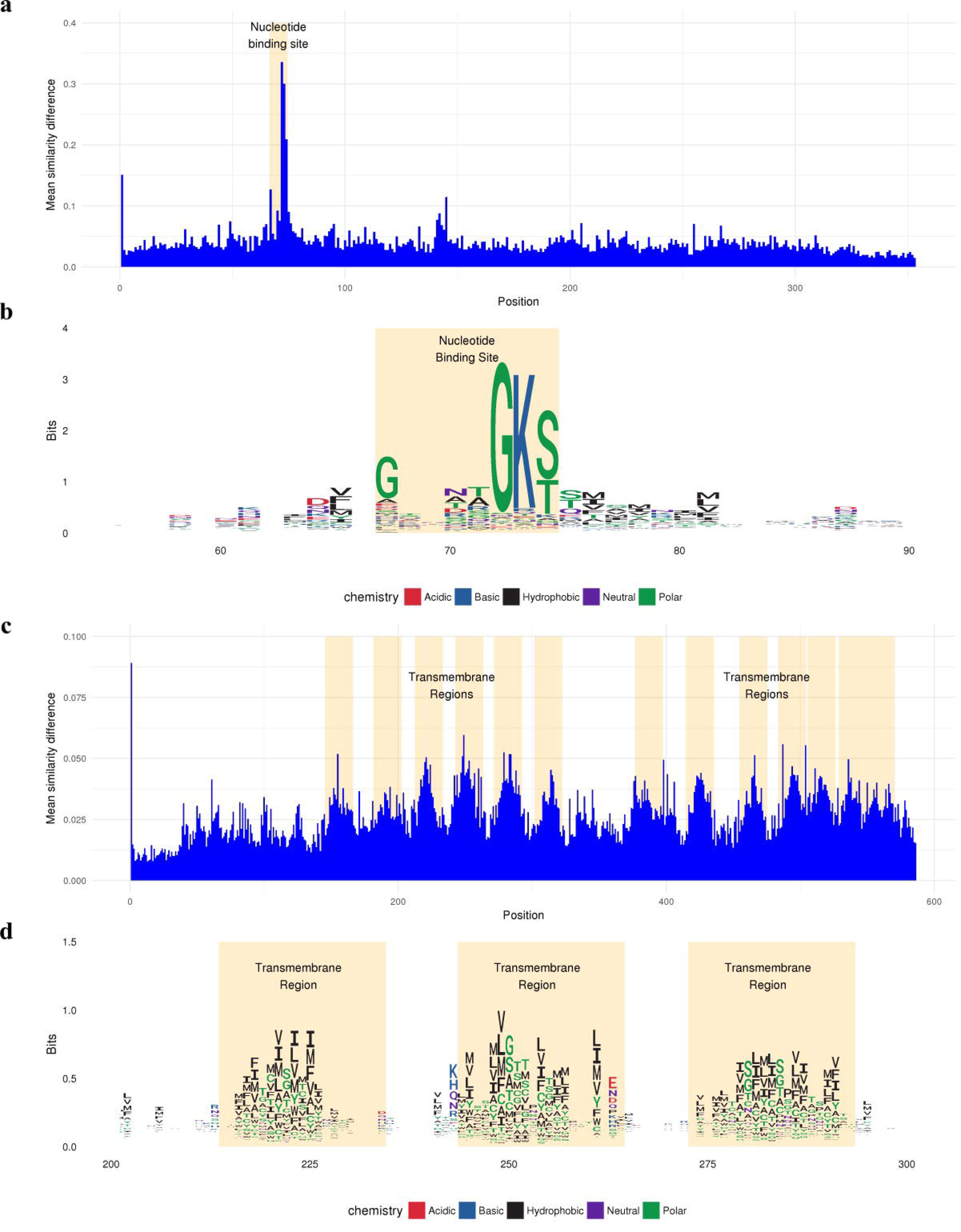
In silico mutagenesis. Two in silico full scans were performed with the D-SPACE model. Each residue in the protein was individually changed to each of the other 19 amino acids and the embedding was recalculated. The functional impact of each substitution is measured as the similarity distance from the original embedding. (**a**) The average functional impact of a substitution at each amino acid for RecA (UniProt P0A7G6). (**b**) A sequence logo view of the peak modifications for RecA. The tallest letters represent amino acids predicted by D-SPACE to be the most critical for maintaining protein function, and recapitulate the well-known NTP binding P-loop motif ^37^. Accordingly, UniProt annotates positions 67-74 as the nucleotide binding site. (**c**) The average functional impact of a substitution at each amino acid for Tpo1 (UniProt Q07824). The UniProt annotated transmembrane regions match the D-SPACE functional impact predictions. (**d**) A sequence logo view of some transmembrane regions for Tpo1.

## Discussion

Deep learning models such as D-SPACE provide dramatic capability improvements to protein biology, including rapid annotation, search, and new ways of generalizing knowledge beyond single labels. The increase in speed allows for the rapid annotation of large databases such as UniProt or GenBank. This has long been a challenge for groups like JGI, NCBI and EBI who maintain large, ever-increasing sequence databases that benefit from periodic reannotation with new and updated models as they become available. Additionally, it is now possible to generate extremely large and complex metagenomes. In our own work (not reported here) we recently assembled a 47 GB metagenome and identified more than 80 million protein-coding genes. To annotate them with a comprehensive conventional pipeline might take months and cost several hundred thousand dollars in compute resources. With D-SPACE we performed the annotations in hours and for less than $1000.

D-SPACE also serves as a prototype for revolutionizing molecular biology by integrating knowledge from across the field in a comprehensive and synergistic manner. Without the bias of hand-curated rules, the model finds meaningful patterns such as active sites and transmembrane regions *ab initio*. We foresee a future where a model such as D-SPACE serves as an anchor for bringing vast amounts of biological information into a single understanding. For starters, D-SPACE can be extended to include additional knowledge about proteins, including enzyme kinetics, thermal stability, three-dimensional structure, etc. The model could also extend upstream to DNA, providing information for coding and non-coding regions alike. With some creative deep learning architectures, almost any conceivable experiment can contribute to the whole, including interaction networks, gene expression, epigenetics, phenotypic effects, drug binding, and clinical outcomes. Each extension of the core model not only adds to its basic utility, but also provides synergistic information relevant to each specific domain of study. For example, the same patterns found useful for annotating protein function are likely to be useful for interpreting drug binding and vice versa.

The rapid advancement in the field of artificial intelligence will likely bring even more powerful capabilities. Reinforcement learning and generative models are proving to be extraordinarily powerful for other fields such as robotics and computer vision^38^. We are approaching a time when artificial intelligences can synthesize imagery, music, and even human speech from a list of specifications with uncanny accuracy. Applying the same approaches to a framework such as D-SPACE could give biologists unprecedented power to engineer proteins for specific tasks or even to create proteins with completely novel functionality.

## METHODS

### Data processing

UniProt data from the February 2018 release was downloaded and parsed into JSON lines format containing a single protein per line. To deduplicate the dataset, records were removed if they were not representative members of a UniRef100 cluster. OrthoDB assignments were extended from a small subset of proteins to the entire dataset with the use of Diamond^39^. To assign a cluster, we required the top hit to have a minimum score of 40 and a minimum of 35% coverage for both the query and subject. Keywords were extracted from the protein description field based on a custom string processing function which attempted to account for a myriad of oddities (ex. hypothetical proteins). Gene Names were also standardized with a custom function available in the code repository. A total of 91 million proteins were randomly split into training (80%), validation (10%), and test (10%) groups. These were shuffled to ensure that records were processed in a random order.

Records were filtered from model training if the sequence contained non-canonical amino acids (ex. ‘X’), was longer than 2000 amino acids, or contained no multi-hot annotations other than keywords. Roughly 3% of records were filtered this way.

### Model construction

The D-SPACE model was built using TensorFlow with Keras (www.tensorflow.org). It consists of a convolutional sequence encoder followed by affine layers to produce the 256-value embedding. This embedding is attached to a decoder for each output task (**Supplemental Fig. 1**). An affine autoencoder task was also added to produce a three-dimensional representation of the embedding layer. The model was trained with the NADAM optimizer with an initial learning rate of 0.001^40^. We chose loss weights for each output to ensure each had a meaningful impact on the overall loss. The model was trained using a Tesla P100 GPU for nine days, until the validation loss no longer improved.

The specification of this architecture requires a number of hyperparameters, including the number and types of layers, the associated layer parameters, the choice of optimizer, the initial learning rate, and the loss weights. These hyperparameters were chosen with a combination of intuition from existing literature and targeted hyperparameter scans. For each trial run, the model loss for training and validation samples were carefully monitored to prevent overfitting. The test data was never used during this procedure, so it could be used to accurately assess the final model’s performance.

### Determination of ‘optimal’ task thresholds

For each task, we determined the scoring threshold (>0.1) at which the F1 score was maximal in the validation dataset.

### Protein similarity metric

The embedding similarity metric is based on the Euclidean distance between two embedding vectors e_1_ and e_2_.

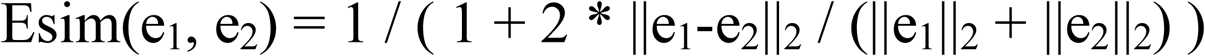

### Defining a ‘non-trivial’ annotation description

We defined an annotation as being non-trivial if it was not ‘None’ and did not contain any of the following text, “hypothetical”, “unknown function”, “uncharacterized protein”, “uncharacterized conserved protein”, “uncharacterized protein conserved in bacteria”, “putative”.

### Code availability

All code necessary for building a D-SPACE model and running inference is available on GitHub at https://github.com/syntheticgenomics/sgidspace. Several utility scripts are also provided to help with constructing the training dataset. This code is available under a GNU Affero General Public License v3.0.

## Acknowledgements

We would like to thank Todd Peterson and Amir Khosrowshahi for supporting the SGI/Intel collaboration.

## Author contributions

A.S.S., T.H.R, A.K.B, led and organized the project. A.S.S, A.R.G, M.E.S, S.A.B processed the training and evaluation datasets. G.J.H., Z.R.D, A.S.S, M.E.S, A.R.G, J.M.K, H.E., Y.L, developed the model architecture, code, and training parameters. A.S.S., G.J.H., A.R.G, M.C.L, S.A.B produced additional analyses. G.J.H, A.S.S., T.H.R, S.A.B, A.R.G wrote the paper. J.R.E, M.E.S, A.S.S, G.J.H, produced the associated website.

## Competing interests

This work was funded and developed by Synthetic Genomics, Inc. and Intel (work was initiated at Nervana Systems Inc.).

**Supplemental Figure 1** D-SPACE network architecture

A visualization of the deep learning model architecture at the heart of D-SPACE.

**Supplementary Figure 2** D-SPACE class label performance

For each multi-hot task label, the F1 statistic observed in our test set is shown in relation to the number of protein records for the label that were present in the training set. Trend lines were fit with a weighted second-degree LOWESS and are displayed with a 95% confidence interval.

## References

1. Stephens, Z. D. et al. Big Data: Astronomical or Genomical? PLOS Biol. 13, e1002195 (2015).

2. Benson, D. A. et al. GenBank. Nucleic Acids Res. 41, D36–D42 (2012).

3. UniProt Consortium, T. UniProt: the universal protein knowledgebase. Nucleic Acids Res. 46, 2699–2699 (2018).

4. Hutchison, C. A. et al. Design and synthesis of a minimal bacterial genome. Science (80-.). (2016). doi:10.1126/science.aad6253

5. Radivojac, P. et al. A large-scale evaluation of computational protein function prediction. Nat. Methods 10, 221–227 (2013).

6. Altschul, S. F., Gish, W., Miller, W., Myers, E. W. & Lipman, D. J. Basic local alignment search tool. J. Mol. Biol. 215, 403–410 (1990).

7. Eddy, S. R. Accelerated profile HMM searches. PLoS Comput. Biol. (2011). doi:10.1371/journal.pcbi.1002195

8. Punta, M. et al. The Pfam protein families database. Nucleic Acids Res. 40, D290–D301 (2012).

9. Haft, D. H., Selengut, J. D. & White, O. The TIGRFAMs database of protein families. Nucleic Acids Res. 31, 371–3 (2003).

10. Pandit, S. B. et al. SUPFAM: A database of sequence superfamilies of protein domains. BMC Bioinformatics 5, 28 (2004).

11. Lees, J., Yeats, C., Redfern, O., Clegg, A. & Orengo, C. Gene3D: merging structure and function for a Thousand genomes. Nucleic Acids Res. 38, D296–D300 (2010).

12. Schultz, J., Milpetz, F., Bork, P. & Ponting, C. P. SMART, a simple modular architecture research tool: identification of signaling domains. Proc. Natl. Acad. Sci. U. S. A. 95, 5857–64 (1998).

13. Mulder, N. J. et al. InterPro: an integrated documentation resource for protein families, domains and functional sites. Brief. Bioinform. 3, 225–35 (2002).

14. Sigrist, C. J. A. et al. PROSITE, a protein domain database for functional characterization and annotation. Nucleic Acids Res. 38, D161–6 (2010).

15. Jurtz, V. I. et al. An introduction to deep learning on biological sequence data: examples and solutions. Bioinformatics 33, 3685–3690 (2017).

16. Kulmanov, M., Khan, M. A., Hoehndorf, R. & Wren, J. DeepGO: predicting protein functions from sequence and interactions using a deep ontology-aware classifier. Bioinformatics 34, 660–668 (2018).

17. Jensen, L. J., Skovgaard, M. & Brunak, S. Prediction of novel archaeal enzymes from sequence-derived features. Protein Sci. 11, 2894–8 (2002).

18. Lee, B. J., Shin, M. S., Oh, Y. J., Oh, H. S. & Ryu, K. H. Identification of protein functions using a machine-learning approach based on sequence-derived properties. Proteome Sci. 7, 27 (2009).

19. Arango-Argoty, G. et al. DeepARG: a deep learning approach for predicting antibiotic resistance genes from metagenomic data. Microbiome 6, 23 (2018).

20. Liu, X. L. Deep Recurrent Neural Network for Protein Function Prediction from Sequence. *arXiv* (2017). doi:10.1101/103994

21. Fa, R., Cozzetto, D., Wan, C. & Jones, D. T. Predicting human protein function with multi-task deep neural networks. PLoS One 13, e0198216 (2018).

22. Nauman, M., Rehman, H. U., Politano, G. & Benso, A. Beyond Homology Transfer: Deep Learning for Automated Annotation of Proteins. *bioRxiv* 168120 (2017). doi:10.1101/168120

23. Qi, Y., Oja, M., Weston, J. & Noble, W. S. A unified multitask architecture for predicting local protein properties. PLoS One (2012). doi:10.1371/journal.pone.0032235

24. Suzek, B. E. et al. UniRef clusters: a comprehensive and scalable alternative for improving sequence similarity searches. Bioinformatics 31, 926–32 (2015).

25. Melvin, I., Weston, J., Noble, W. S. & Leslie, C. Detecting remote evolutionary relationships among proteins by large-scale semantic embedding. PLoS Comput. Biol. (2011). doi:10.1371/journal.pcbi.1001047

26. Loh, P. R., Baym, M. & Berger, B. Compressive genomics. Nature Biotechnology (2012). doi:10.1038/nbt.2241

27. Daniels, N. M. et al. Compressive genomics for protein databases. in Bioinformatics (2013). doi:10.1093/bioinformatics/btt214

28. Yu, Y. W., Daniels, N. M., Danko, D. C. & Berger, B. Entropy-Scaling Search of Massive Biological Data. Cell Syst. 1, 130–140 (2015).

29. Ruder, S. An Overview of Multi-Task Learning in Deep Neural Networks. *arXiv* (2017). doi:10.1109/CVPR.2015.7299170

30. Andoni, A., Indyk, P., Laarhoven, T., Razenshteyn, I. & Schmidt, L. Practical and Optimal LSH for Angular Distance. Adv. Neural Inf. Process. Syst. 28 (2015).

31. Johnson, J., Douze, M. & Jégou, H. Billion-scale similarity search with GPUs. arXiv 1702.08734 (2017).

32. Rost, B. Twilight zone of protein sequence alignments. Protein Eng. Des. Sel. (1999). doi:10.1093/protein/12.2.85

33. Saripella, G. V., Sonnhammer, E. L. L. & Forslund, K. Benchmarking the next generation of homology inference tools. Bioinformatics 32, 2636–2641 (2016).

34. Altschul, S. F. et al. Gapped BLAST and PSI-BLAST: a new generation of protein database search programs. Nucleic Acids Res. 25, 3389–402 (1997).

35. Johnson, L. S., Eddy, S. R. & Portugaly, E. Hidden Markov model speed heuristic and iterative HMM search procedure. BMC Bioinformatics 11, 431 (2010).

36. Price, M. N. et al. Mutant phenotypes for thousands of bacterial genes of unknown function. Nature 557, 503–509 (2018).

37. Saraste, M., Sibbald, P. R. & Wittinghofer, A. The P-loop--a common motif in ATP- and GTP-binding proteins. Trends Biochem. Sci. 15, 430–4 (1990).

38. Karras, T., Aila, T., Laine, S. & Lehtinen, J. Progressive Growing of GANs for Improved Quality, Stability, and Variation. arXiv 1710.10196 (2017).

39. Buchfink, B., Xie, C. & Huson, D. H. Fast and sensitive protein alignment using DIAMOND. Nat. Methods 12, 59–60 (2015).

40. Dozat, T. Incorporating nesterov momentum into adam. ICLR 2016 (2016).

